# Enhancing the Robustness of OPLS Modelling in Small Cohorts by Leveraging Permutation Analysis Prior to Variable Selection

**DOI:** 10.1101/2024.03.18.585475

**Authors:** Marika Ström, Nicole Wagner, Iryna Kolosenko, Åsa M. Wheelock

## Abstract

The R-workflow ropls-ViPerSNet (**R o**rthogonal **p**rojections of **l**atent **s**tructures with **V**ar**i**able **Per**mutation **S**election and Elastic **Net**) facilitates variable selection, model optimization and significance testing using permutations of OPLS-DA models, with the scaled loadings (p[corr]) as the main metric of significance cutoff. Permutations including (over) the variable selection procedure, prior to (pre-), as well as post variable selection are performed. The resulting p-values for the correlation of the model (R^2^) and the cross-validated correlation of the model (Q^2^) pre-, post- and over-variable selection are provided as additional model statistics. These model statistics are useful for determining the true significance level of OPLS models, which otherwise have proven difficult to assess particularly for small sample sizes. Furthermore, a means for estimating the background noise level based on permuted false positive rates of R^2^ and Q^2^ is proposed. This novel metric is then used to calculate an adjusted Q^2^ value. Using a publicly available metabolomics dataset, the advantage of performing permutations over variable selection was demonstrated for small sample sizes. Iteratively reducing the sample sizes resulted in overinflated models with increasing R^2^ and Q^2^ and permutations post variable selection indicated falsely significant models. In contrast, the adjusted Q^2^ was marginally affected by sample size, and represents a robust estimate of model predictability, and permutations over variable selection showed true significance of the models. An additional Elastic Net variable selection option is included in the workflow for variable selection by coefficient value penalization using an iterative approach to reduce noise while avoiding overfitting.

## Background

In clinical studies, particularly those involving invasive sampling, multivariate statistical modeling is key to characterizing disease mechanisms. Small sample sizes combined with a multitude of analytes in omics studies increase the risk of false discoveries [2].

Partial least squares (PLS) is a multivariate modelling method [3] used for correlating two data blocks. When one block (generally the Y-block) is set to a single variable, PLS can be used as a supervised method to evaluate group separation in a multivariate fashion, as well as to extract features driving the separation. Orthogonal projections to latent structures (OPLS) [4] is a development of PLS that rotates the principal components in order to separate the variance *within* the groups of interest (orthogonal components) from the variance *between* the groups of interest (predictive components), and thereby structure the model to better facilitate the extraction of features driving the separation between the groups. Similar to PLS, the y-block needs to be reduced to variables that best determine group separation to facilitate efficient execution of supervised modeling.

OPLS is an efficient tool for omics dataset analysis [5]. The model statistics provided by the original algorithms in the SIMCA software [6] are primarily designed to provide significance of group separation through cross-validated ANOVA (CV-ANOVA), and the predictive power of group separation through the cross validated correlation of the original Y-block with the model (Q^2^Y, a.k.a. Q^2^). The resulting OPLS model can help reduce the large number of variables in an omics dataset to a subset of key features that drive group separation, which may be of interest as potential biomarkers or molecular targets. This process, known as variable selection or feature selection, is often conducted using algorithms such as random forest, support vector machine, or an alpha-cutoff based on t-statistics corrected for multiple testing.

In OPLS, two main variable selection metrics are commonly used: Variable Importance in the Projection (VIP) and the scaled loadings (p[corr]) [5]. VIP ranks variables based on their contribution to group separation, with 1.0 representing the average contribution. The relative nature of VIP and its dependence on the number of variables make it suboptimal for variable selection, particularly when comparing variable contribution between models. In this context, p(corr) is a more robust metric given that it is related to the loadings of the predictive variable and is scaled as a correlation coefficient ranging from -1 to 1, thereby facilitating direct comparison between models. As such, using p(corr) as the primary metric for variable selection in OPLS is proposed.

To evaluate the performance of models, R^2^Y (hereafter R^2^) is used. R^2^ is the established model statistic that describes the variation of the dataset explained by the model. Q^2^Y (hereafter Q^2^) is also considered. Q^2^ describes the predicted R^2^ assessed by cross validation and is a more robust measure of the model performance [7].

OPLS models are prone to overfitting [8], which can be assessed by adding additional principal components to a model. This increases R^2^ and can increase group separation but at the same time the predictability Q^2^ decreases indicating over-fitting of the model. Reducing the ratio between the number of samples and features leads to greater separation, potentially resulting in perfect segregation of a dataset when the ratio is low. This threshold has been estimated to be 1:2 or lower [9]. Also, during variable selection R^2^ and Q^2^ increase, and the extent of the increase is partly due to the overinflation of the model statistics. CV-ANOVA is a common method for OPLS testing; in this context accuracy is defined as the proportion of the subject group correctly predicted by the model.

One way to test the significance of model statistics is by permutation tests [10]. In this approach, sample group labels are scrambled prior to modeling the dataset. The procedure is iterated, and the results represent model statistics unaffected by chance. Permutation tests, such as SIMCA software (Sartorius) [6], have been implemented in tools for multivariate analysis and in the R package ropls. This procedure is commonly performed either pre-variable selection (by including all variables in the full dataset to calculate the significance of R^2^ and Q^2^), or post-variable selection (by including only the final selected variables of interest in the permutation test). While the latter approach assures that the selected variables will not produce a significant separation of groups of subjects randomly picked from cohort, it brings the caveat of only controls for the variables of interest being considered and does not assess alterations represented by random variation.

A more robust method to assess significance involves permutation testing with variable selection, also known as permutation test over variable selection. In this procedure, permutations are first conducted similarly to standard permutation testing. Then, variable selection is applied to each permuted dataset before fitting a model to the permuted data. This approach, which incorporates variable selection into permutations, has been explored in OPLS models [11] and has been integrated into the MUVR package for PLS models [12].

To estimate the significance of the model, the model statistics of permuted models to those of the unpermuted model under examination are compared. This is achieved by calculating the ratio between permuted models that exhibit statistics above and below those of the unpermuted reference model, yielding a nonparametric p-value. This approach, implemented in the ropls package, provides p-values for R^2^ and Q^2^.

The objective of this study was to develop an R workflow for OPLS modeling, termed ropls-ViPerSNet, that incorporates variable selection and permutations both before and after variable selection . Additionally, a significance level for the predictive power of Q^2^ in models post-variable selection is established by comparing it to the estimated overinflation caused by reducing the number of variables. This overinflation serves as a threshold for Q^2^, with models achieving a Q^2^ above this threshold considered statistically significant. Subsequently, an adjusted Q^2^ was calculated by subtracting the estimated threshold, resulting in a significance cutoff of 0.

To mitigate overfitting in datasets at risk due to limited sample sizes, regularization techniques like Lasso, Ridge, and Elastic Net regression are implemented using the glmnet model [13] training framework within the caret R package [14]. These methods address overfitting and multicollinearity by adding penalty terms to the loss function, which minimizes the error between predicted and actual values. Ridge regression (L2 penalty) shrinks coefficients towards zero without eliminating predictors, while Lasso regression (L1 penalty) promotes sparsity by setting irrelevant coefficients to zero, thereby performing variable selection. Elastic Net combines these penalties, balancing Ridge and Lasso via the mixing parameter alpha (alpha=1 for Lasso, alpha=0 for Ridge), making it effective for datasets with correlated predictors or more variables than samples. The degree of regularization is controlled by the lambda parameter. Subsequent variable selection informs an OPLS model, whose performance is evaluated using Q² and R² metrics. It should be noted that this approach requires larger computational capacity.

In this study, the script is used to examine the impact of decreasing sample sizes on R^2^, Q^2^, overinflation, and the proposed adjusted Q^2^ using a publicly available metabolomics dataset. Furthermore, whether permutations pre-, post-, and over variable selection could distinguish between a significant model and a model generated from random data at various sample sizes was investigated. Given that OPLS models are prone to overfitting, particularly after variable selection, the development of a tool to identify and avoid publishing insignificant models is crucial. Such a tool could help address the well-established issue of irreproducibility in research.

## Methods

### The ropls-ViPerSNet Workflow

The ropls-ViPerSNet workflow is based on R and designed to streamline semi-automated pairwise group comparisons within a dataset using OPLS modeling, based on primary group assignment (e.g., treatment vs. placebo groups). The workflow offers the flexibility to stratify the analysis based on additional variables of interest (e.g., gender, smoking status). To ensure robust and reliable model optimization thorough significance testing, the workflow builds on permutations (5 Modelling strategies) and the option for using regression analysis for variable penalization, which incorporate variable selection.

OPLS analysis is conducted using the Bioconductor R package ropls [15, 16]. Additionally, for generating HTML reports with plots, the package relies on several dependencies including Rmarkdown [17, 18], tools [19] ggplot2 [20], ggrepel [21], kableExtra [22], gridExtra [23], ggpubr [24], matrixStats [25], stringr [26], tryCatchLog [27], devtools [28], DescTools [29], precrec [30], pROC [31], rstatix [32], glmnet [13] and caret [14].

The ropls-ViPerSNet R workflow is available at https://github.com/pulmonomics-lab/ropls_vipersnet.

### Workflow

The workflow of ropls-ViPerSNet entails running a main file (oplspvs.R) that executes the opls function (Figure 1). Users can customize a subset of settings through two configuration files:

1. The Configure_Get_Started.R file contains basic parameter settings, including information about the data matrix (e.g., variable names, preprocessing details) and observations (e.g., sample IDs, primary group assignments). Additionally, it specifies the metadata to be used for stratification in OPLS analyses. This file also allows users to define the number of permutations, the significance level for the p(corr) cutoff, and the order of groups for directional interpretation in OPLS score plots (e.g., healthy vs. diagnosis or diagnosis vs. healthy). Users can also specify which pairwise comparisons to run, with the default being all groups in the sample ID file.
2. The Configure_Advanced.R file enables users to customize variable selection settings, set the maximum number of allowed orthogonals, define criteria for selecting the best performing model, and adjust the length and format of variable names displayed in OPLS loadings plots. Users can select modelling strategies (default: 0-5) and choose whether to generate both comparisons and summary output files. This file also offers users the option of running the Elastic Net variable selection approach and adjust alpha values and lambda thresholds for this approach.

**Figure 1.**
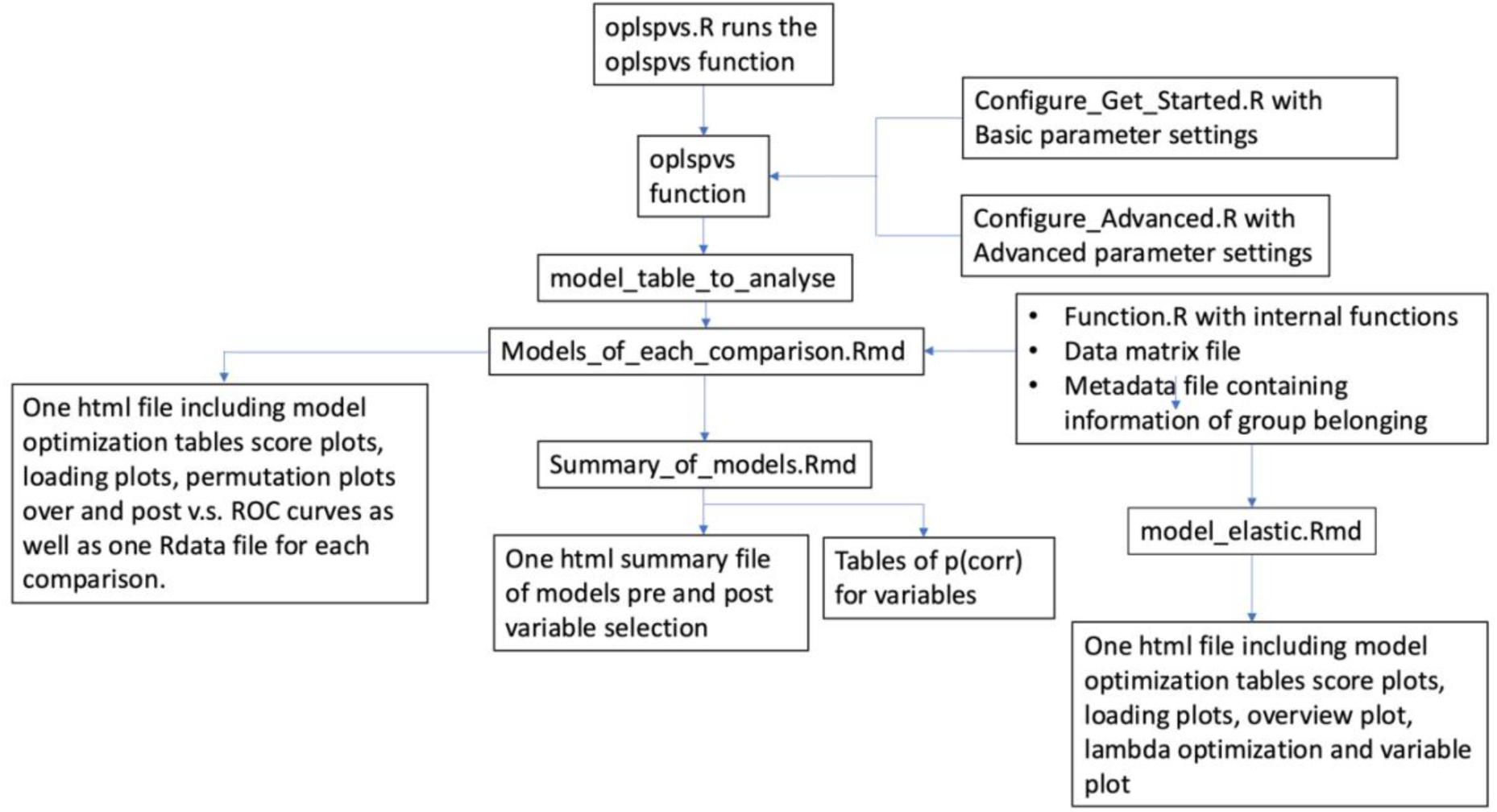
Schematic of the ropls-ViPerSNet workflow.

The ropls-ViPerSNet script first generates a table (model_table_to_analyse) containing information about each pairwise comparison to be performed. Then, it passes parameters from the model_table_to_analyse and the two configure files to the Rmarkdown files. The data matrix, metadata file, and functions from roplspvs_Functions.R are incorporated into the Rmarkdown files to conduct the modelling, resulting in an Rdata file and an HTML output file for each comparison. The HTML output includes score plots, loading plots, permutation plots before and after variable selection, and Receiver Operating Characteristic (ROC) curves. Additionally, HTML files summarizing model statistics for all pairwise comparisons are generated, along with variable lists displaying associated p(corr) values for variables selected by the respective modeling strategies. A separate HTML file with the same information will be generated if users choose to also run Elastic Net regression-based variable selection. Figure 1 provides an overview of the input and output files of ropls-ViPerSNet.

### Preprocessing of Dataset

The dataset undergoes preprocessing as outlined in Figure S1. Preprocessing is iteratively conducted for each pairwise comparison to optimize the variables considered. Options include replacing zero values with NA or the lower limit of detection (LLD), followed by optionally replacing NA with LLD. Users can manually set the LLD or estimate it as 1/3 of the minimum value in the dataset. Additionally, data filtering removes variables based on a user-defined missing value tolerance level for each group in the comparison.

### OPLS and PCA Modeling

The ropls package facilitates initial PCA and OPLS modeling [15]. By default, OPLS employs NIPALS (Nonlinear Iterative Partial Least Squares) to handle missing data, if NAs are present in the dataset. Mean centering and scaling to unit variance (UV) are applied, enabling variables with low abundance or amplitude to contribute to the model.

The process begins with the generation of PCA models pre-variable selection for each comparison. This encompasses creating PCA models that integrate all observations in the comparison, as well as PCA models for each group individually, aiding in robust outlier identification. Subsequently, OPLS models are constructed for each comparison, both pre-variable selection (encompassing all variables meeting the preprocessing QC criteria) and post-variable selection (comprising solely variables selected by the respective modeling strategy, as elaborated in the section below). Model evaluation entails reporting R^2^, Q^2^, and RMSE model statistics, alongside significance assessment through permutations.

### Variable Selection

The first approach includes five OPLS modeling strategies include feature selection, also termed variable selection, operates using five distinct modeling strategies applying different stringency levels. The default variable selection procedures in all 5 model strategies hinge on p(corr), with the possibility of integrating Variable Importance in Projection (VIP) as an additional variable selection metric. Different p(corr) cutoffs for variable selection are applied across the five strategies. The determination of the p(corr) cutoff primarily stems from the iterative optimization of the predictive power, Q^2^, of the model post-variable selection. This iterative process aims to thwart overfitting, thus maintaining the maximum predictive power of the model after variable selection. Figure 2 provides an overview of the available modeling strategies.

**Figure 2.**
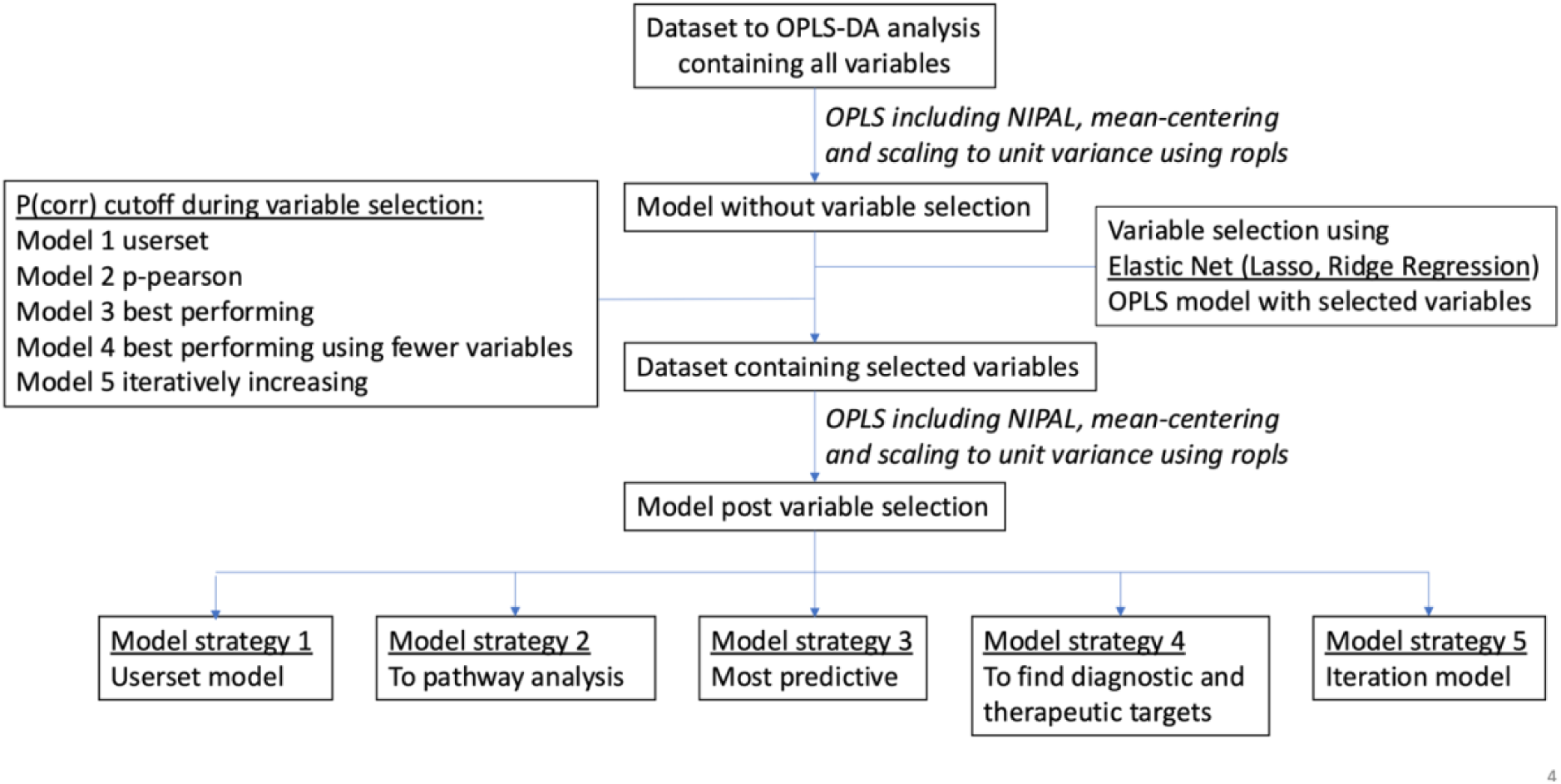
Overview of the five different Modelling strategies available in the ropls-ViPerSNET package. The p(corr) cutoff used in the respective modelling strategies are indicated in the text box to the left.

The second optional approach involves Lasso, Ridge, or Elastic Net regression, requiring two hyperparameters: lambda and alpha. For Elastic Net, alpha (ranging from 0 for Ridge to 1 for Lasso) is user-defined, while the optimal lambda is selected via cross-validation. A sequence of lambda values, with minimum and maximum values set by the user, is evaluated by training models on all but one fold of the data and testing on the held-out fold. This is repeated across all folds, and the mean cross-validated loss (e.g., negative log-likelihood for classification) is calculated for each lambda. The lambda minimizing the loss is chosen for the final model, with the largest lambda within one standard error of this minimum selected to encourage parsimony. Variable selection is performed using the optimal lambda, with non-zero coefficients indicating the selected variables.

### Modeling Strategies

In Modeling Strategy 1, users have the option to set the p(corr) cutoff, offering flexibility to adjust model settings or evaluate models optimized in other software using the permutation analyses in the ropls-ViPerSNET script.

In Modeling Strategy 2, the p(corr) cutoff is determined by incorporating all variables with significant correlation to the score, established by a user-defined p-value corresponding to the Pearson’s correlation coefficient for the group size. This cutoff also serves as the default for Modeling Strategy 1 if no other cutoff is specified.

Modeling Strategy 3 is recommended for users aiming to identify the best performing model with high Q^2^, minimal difference between R^2^ and Q^2^, and low permutation p-values (p[R2_perm_post_vs] and p[Q2_perm_post_vs]). This strategy prioritizes predictive power while avoiding overfitting and selecting models not significantly better than random. Users can adjust the balance between these criteria using the variable prefered_pR2_and_pQ2_permuted_post_vs.

Modeling Strategy 4 minimizes the number of selected variables while preserving predictability. The p(corr) cutoff is adjusted to achieve the best performing model post-variable selection, gradually increasing the cutoff as long as Q^2^ is not reduced by more than 1%. Users can control the reduction in variables by setting the pcorr_diff parameter. Modeling strategies 3 and 4 are summarized in Figure 3.

**Figure 3.**
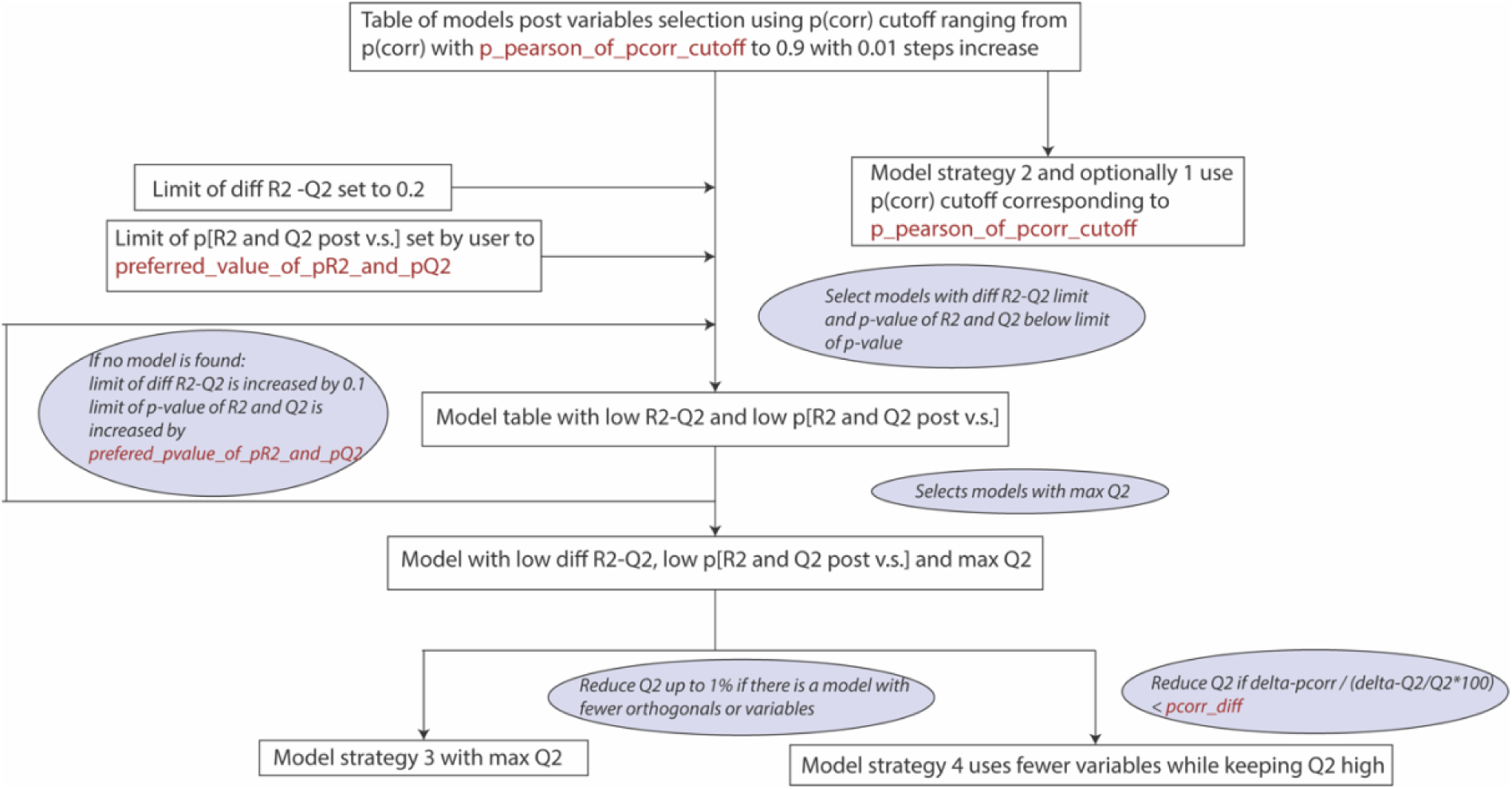
Details for selecting p(corr) cutoff of Modelling strategies 3 and 4. Parameters that are user-defined in ropls-ViPerSNet are marked in red.

Modeling Strategy 5 employs an iterative approach by incrementally increasing the p(corr) cutoff and refining the model, eliminating variables with the least correlation. The cutoff is raised as long as Q^2^ post-variable selection is not reduced by more than 1%.

To facilitate the evaluation of p(corr) cutoff optimization, a plot of Q^2^ versus the number of variables using different cutoffs is generated. This, coupled with a model table, enables users to confirm the selection of the best performing model.

The additional Elastic Strategy, which is separate from the above strategies, focuses on using the power of Lasso, Ridge and Elastic Net Regression, based on user-set hyperparameters, to incrementally increase the penalty parameter lambda which determines the extent of regularization to find the optimal value. A larger lambda imposes stronger regularization, reducing the magnitude of coefficients and potentially setting them to zero (in the case of Lasso and Elastic Net).

### Selection of Number of Orthogonal Variables

The user can define the number of orthogonals for Modeling Strategy 1, with the default set to 0. For Mode**ling** Strategies 2-5, the number of orthogonals is determined using the ropls default method, which adds an orthogonal as long as Q**^2^** increases by 1% in the currently evaluated model. Users can also set the maximum number of orthogonals for both models pre- and post-variable selection. Additionally, the script checks that no strategy produces a better performing model post-variable selection using fewer orthogonals. In the optional regression-based model, orthogonal values between 0 and 2 are tested.

### Permutations

The significance of models using permutations pre-, post-, and over variable selection is assessed, as illustrated in Figure 4.

**Figure 4.**
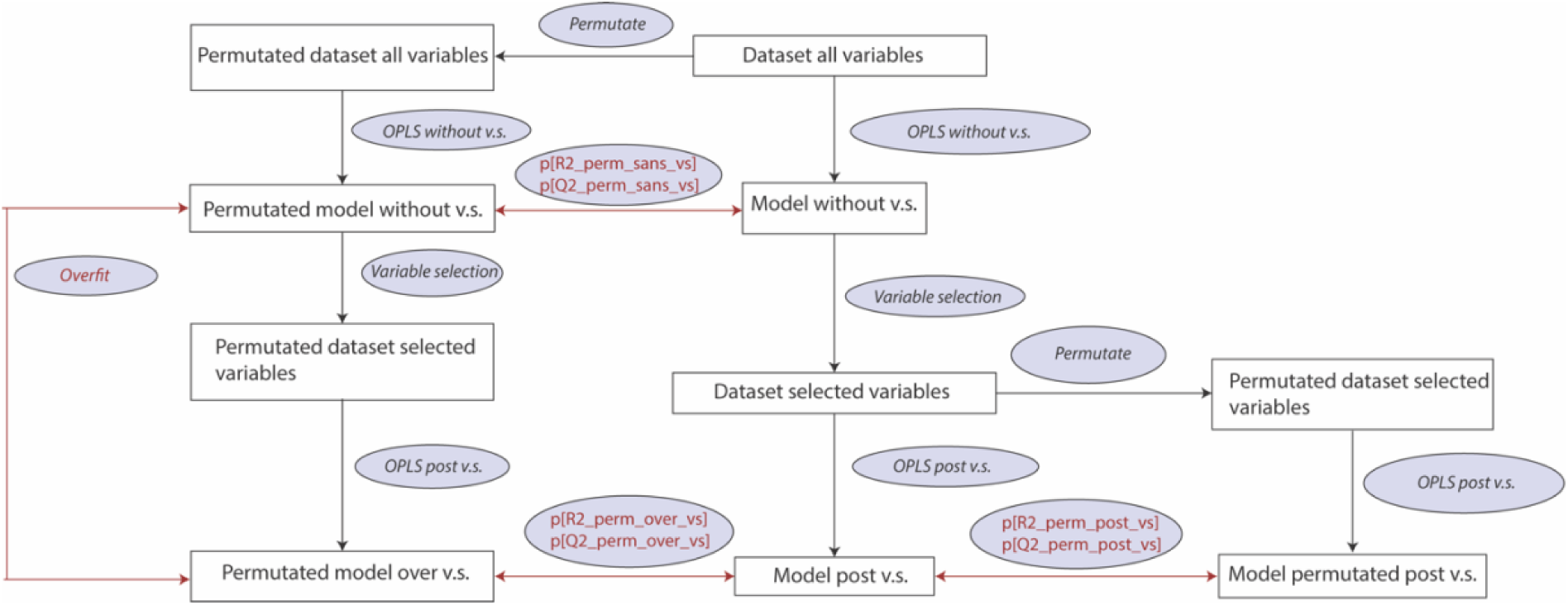
Estimations of the significance level of each model statistics is provided based on permutation test at three levels; prior to variable selection, after variable selection, as well as over variable selection p[R2 and Q2 for permutations sans, post and over variable selection] is estimated by comparing permuted models to unpermuted model under investigation. The comparisons of models for the estimated p-values as well as the overfit are shown in red.

Permutations pre- and post-variable selection involve randomizing the group assignment of subjects using the ropls package. During permutations pre-variable selection, the dataset is modeled by including all variables from preprocessing, against permuted group assignments. In contrast, permutations post-variable selection entail modeling the dataset only with variables selected during variable selection against permuted group assignments. The model statistics R^2^ and Q^2^ from permuted models to those of the original unpermuted models are compared, resulting in p-values for R^2^ and Q^2^ pre-variable selection (p[R2_perm_sans_vs] and p[Q2_perm_sans_vs]) and post-variable selection (p[R2_perm_post_vs] and p[Q2_perm_post_vs]).

Similarly, permutations over variable selection involve modeling the dataset with all variables from preprocessing. Each permuted model undergoes variable selection using the same p(corr) cutoff as the unpermuted model under investigation. The model statistics R^2^ and Q^2^ from permuted models post-variable selection are compared to those of unpermuted models post-variable selection, resulting in p-values for R^2^ and Q^2^ over variable selection (p[R2_perm_over_vs] and p[Q2_perm_over_vs]).

One limitation of permutations over variable selection is that the settings of the unpermuted model are used for permuted models. This may lead to increased significance for p[R2_perm_over_vs] and p[Q2_perm_over_vs] compared to p(corr) being optimized for permuted models as well. To address this, the regression over R^2^ and Q^2^ of permuted models is considered. The R^2^ and Q^2^ of permuted models against the correlation of permuted and unpermuted group assignments is plotted. A significant positive correlation coefficient in the regression indicates a significant model, as higher correlation to the original group assignments should yield higher R^2^ and Q^2^.

The user can set the number of permutations, with a default of 20 permutations for initial analysis, balancing computational time with statistical power. However, increasing the number of permutations is recommended for final analysis.

### Establishing a Statistically Sound Cutoff for Q^2^

The overinflation of OPLS models resulting from variable selection by comparing the Q^2^ of permuted models post-variable selection to the Q^2^ of permuted models pre-variable selection was determined. This difference in Q^2^ represents the overinflation of Q^2^, serves as a robustness threshold. Models with Q^2^ above this threshold are unlikely to have occurred by chance. To account for this overinflation, an adjusted Q^2^ was calculated by subtracting the estimated overinflation from the Q^2^ of the model post-variable selection. Consequently, models with an adjusted Q^2^ above zero can be considered statistically significant and not likely to have occurred by random chance.

### Figures

The ropls package is used to generate scores plots, as well as loading, prediction, diagnostic, and outlier plots, along with plots of permutations without and after variable selection.

Plots generated by ropls-ViPerSNet include score plots of predictive components displayed as boxplots, and p(corr) plots illustrating variables driving model separation. These plots show all variables, and if the number exceeds 50, the 50 most influential variables are displayed. Model optimization is depicted by comparing the number of variables to Q^2^ post-variable selection. Additionally, plots of permutations over variable selection are generated, as described in the section above, using ropls-ViPerSNet.

After model creation and HTML file generation, Shared and Unique Structures (SUS) plots can be used for comparison. SUS plots are correlation plots of p(corr) for each variable in two models [33]. To incorporate unique variables from each model, new models containing all variables from both models are created, with p(corr) from optimized models displayed in the resulting SUS plot. SUS plots aid in identifying variables with similar or different effects when comparing groups, offering insights into potential interactions or confounders in the comparison.

### Qualitative Variables

Qualitative data in the dataset is transformed into dummy variables following the same procedure as in SIMCA workflow, with the adjustment that all dummy variables are included in the modeling also in the case of redundancy. This approach offers the advantage of demonstrating how all settings affect the outcome, despite potentially leading to slight overfitting. Including qualitative variables in the analysis is beneficial for creating models using clinical data.

### Cohort and Dataset

The ropls-ViPerSNET workflow was tested using the untargeted metabolomics dataset MTBLS136 [34], which originates from the Cancer Prevention II Nutrition Cohort [35]. This dataset is publicly available at Metabolights (http://www.ebi.ac.uk/metabolights/MTBLS136/files). It consists of blood metabolome profiles generated on the Metabolon platform for studying post-menopausal hormone treatments in women. Among the 1336 women in the cohort, 332 received estrogen therapy alone (E-only), 337 received combined estrogen and progestin treatment (E+P), and 667 women received no hormone replacement therapy (non-users). For analysis, the women were stratified into age groups of 55 and under, 56-60, 61-65, 66-70, 71-75, 76-80, and 81-85. Zero values in the dataset were replaced with NA, followed by filtering to allow for 25% missing values, and log-transformation.

### Comparing Hormone Users to Non-users Using ropls-ViPerSNet

OPLS modeling using ropls-ViPerSNet was demonstrated with the aforementioned dataset. A maximum of 5 orthogonals was set for both models pre- and post-variable selection to mitigate the risk of overfitting caused by excessive orthogonals. The variable ’prefered_pR2_and_pQ2_permuted_post_vs’ was set to 0.05 to balance between maintaining low p-values for p[R2 and Q2 post v.s.] and minimizing Δ(R^2^-Q^2^), while maximizing Q^2^. Additionally, ’Pcorr_diff’ was set to 0.01 to limit the number of variables in Modeling strategy 4.

For the Elastic Net Regression approach, alpha was set to 0.5, balancing Ridge and Lasso Regression. For Lambda, the range was set to a minimum of 0.001 and maximum of 1, with 1000 intervals. The approach was tested for 0, 1 and 2 orthogonals. The optimal value for Lambda within the range given (0.001 and 1) was found based on the ROC performance of the Elastic Net model (Figure S2).

### Effect of Small Sample Sizes on OPLS Models

The study used the ropls-ViPerSNet package to examine the impact of sample sizes on OPLS models using the previously mentioned dataset. To mitigate unrelated variability based on gender and age, the focus was narrowed to the 61-65 year age group, which is characterized by a relatively large number of participants (n_E+ P_ = 129, n_E-only_= 91, and n_non-users_=121) and homogeneous group nature as confirmed by PCA.

A robust model comparing estrogen plus progestin users to post-menopausal hormone non-users exhibited a Q^2^ of 0.53, deemed significant by permutation tests at all levels (p[Q2_perm_ post_vs] ≤0.01, p[Q2_perm_over_vs] ≤0.01). Conversely, a weaker model with Q^2^ of 0.10 also passed the significance threshold by permutation tests at all levels (p[Q2_perm_sans_vs] ≤0.01, p[Q2_perm_ post_vs] ≤0.01, p[Q2_perm_over_vs] ≤0.01). Additionally, hormone non-users were compared to the same group to verify model insignificance when no difference between groups was expected.

Subsets of subjects from the three models were drawn with consistent sample sizes for each group (spanning from 4 to 80 for each group, 4 to 60 for each non-user group), repeated 12 times for each subset. Modeling strategies 3 and 4 were applied to each subset using ropls-ViPerSNet. R^2^ and Q^2^ for models pre- and post-variable selection were plotted against sample sizes, with models color-coded based on p[R2 and Q2 perm. sans, post, and over v.s.]. Furthermore, the average overinflation and adjusted Q^2^ was determined for each sample size for the strong model.

When smaller subsets of subjects were randomly extracted from each group, models built using Elastic Net exhibited only a small decrease in Q² values. However, the model’s performance still appeared to strongly depend on the number of orthogonal components. Models with zero orthogonal components consistently performed worse than those with one or more orthogonals for this dataset. Detailed results are provided in Supplementary Table 2.

## Results

### Comparing Hormone Users to Non-users Using ropls-ViPerSNet

The effectiveness of the ropls-ViPerSNET workflow was demonstrated using publicly available data investigating blood metabolite composition concerning hormone replacement therapy in post-menopausal women (MTBLS136 [34]). Initially, pairwise comparisons were conducted for all contrasts among the three study groups: estrogen-and-progestin users (E+P), estrogen-only users (E only), and individuals reporting no hormone replacement use (non-users). Analyses for both all age groups (joint) and stratified by age groups using 5-year bins was performed. Age group 81 and above was excluded due to small sample sizes.

Supplementary Table 1 summarizes the model statistics for all pairwise comparisons using Modeling strategy 4. All models across all Modeling strategies demonstrated significance in permutation post-variable selection (p[Q2_perm_post_vs]). Even with the more stringent approach of permutation testing over variable selection (p[Q2_perm_over_vs]), most comparisons remained significant across all Modeling strategies. The exceptions were the E+P versus E-only comparison in the 56-60 age group in Modeling strategy 1 and 2, and the 76-80 age group, which were insignificant using Modeling strategy 1 through 4.

Similarly, p[Q2_perm_sans_vs] indicated significance for all comparisons except the two mentioned above. These two insignificant comparisons also displayed insignificant p-values for correlation of permutations across Modeling strategy 1 through 4. For Modelling strategy 2 through 4, both models pre- and post-variable selection showed low Q^2^ (<0.4) for all age groups comparing E+P versus E-only, while models comparing E+P and E-only versus non-users exhibited high Q^2^ (>0.4). Adjusted Q^2^ was consistently low (<0.1) for all models across all age groups comparing E+P versus E-only.

When applying the Elastic Net approach on the post-variable selection modelling approach. The hyperparameter alpha was set to 0.5 to balance the Ridge and Lasso regression approaches. The hyperparameter lambda was calculated between 0.001 and 1, with 1000 iterations. The OPLS model built with the selected variables resulted in a pQ² (p-value for model prediction) of < 0.001 for all models prior to age stratification. The Q² values were 0.65 for E versus nonusers and 0.70 for E+P versus non-users, with the model efficiency slightly stronger for two orthogonals and slightly weaker for zero orthogonals. Consistent with the initial step (modeling strategies), the efficacy of the model declined when comparing E versus E+P, with Q² values dropping to 0.40 for two orthogonals, 0.32 for one orthogonal, and 0.26 with no orthogonals (Supplementary Table 2).

When models were stratified by age, the Q² values increased across all groups. For one orthogonal, age-stratified Q² values ranged from 0.89 (E+P versus non-users, 56–60 years) to 0.59 (E+P versus E, 61–65 years). Complete results are available in Supplementary Table 2. Figure S3 provides details for the OPLS model using one orthogonal. While the overall model performance is strong, the improved performance is driven by a combination of predictor variables and orthogonal components, highlighting the importance of including orthogonals.

Additionally, as shown in Figure S3, Q² prediction values are consistent with the number of variables used, though prediction performance for the total number of individuals without stratification involves a much higher number of variables. Prior to age stratification, models with two orthogonals provided better predictions than those with one orthogonal. However, following age stratification, the performance of models with one and two orthogonals became comparable, both outperforming models without orthogonals. This trend remained when the age-stratified models were further divided into smaller groups.

The ropls-ViPerSNet workflow generates a plot illustrating the Q^2^ post variable selection against the number of selected variables in the respective model to track iterative model optimization based on p(corr). This plot indicates the p(corr) cutoff for the model, as shown in Figure 5 for optimizing the contrast between E+P and non-users in the 61-65 age group across Modeling strategies 2 through 4. For Modeling strategy 2, a p(corr) cutoff of 0.124 was used, selecting 384 variables with Q^2^=0.56. Modeling strategy 3 employed a p(corr) cutoff of 0.22, selecting 144 variables, resulting in Q^2^=0.58. Finally, Modeling strategy 4 with a p(corr) cutoff of 0.44 reduced the selected variables to 14, yielding Q^2^=0.53.

**Figure 5.**
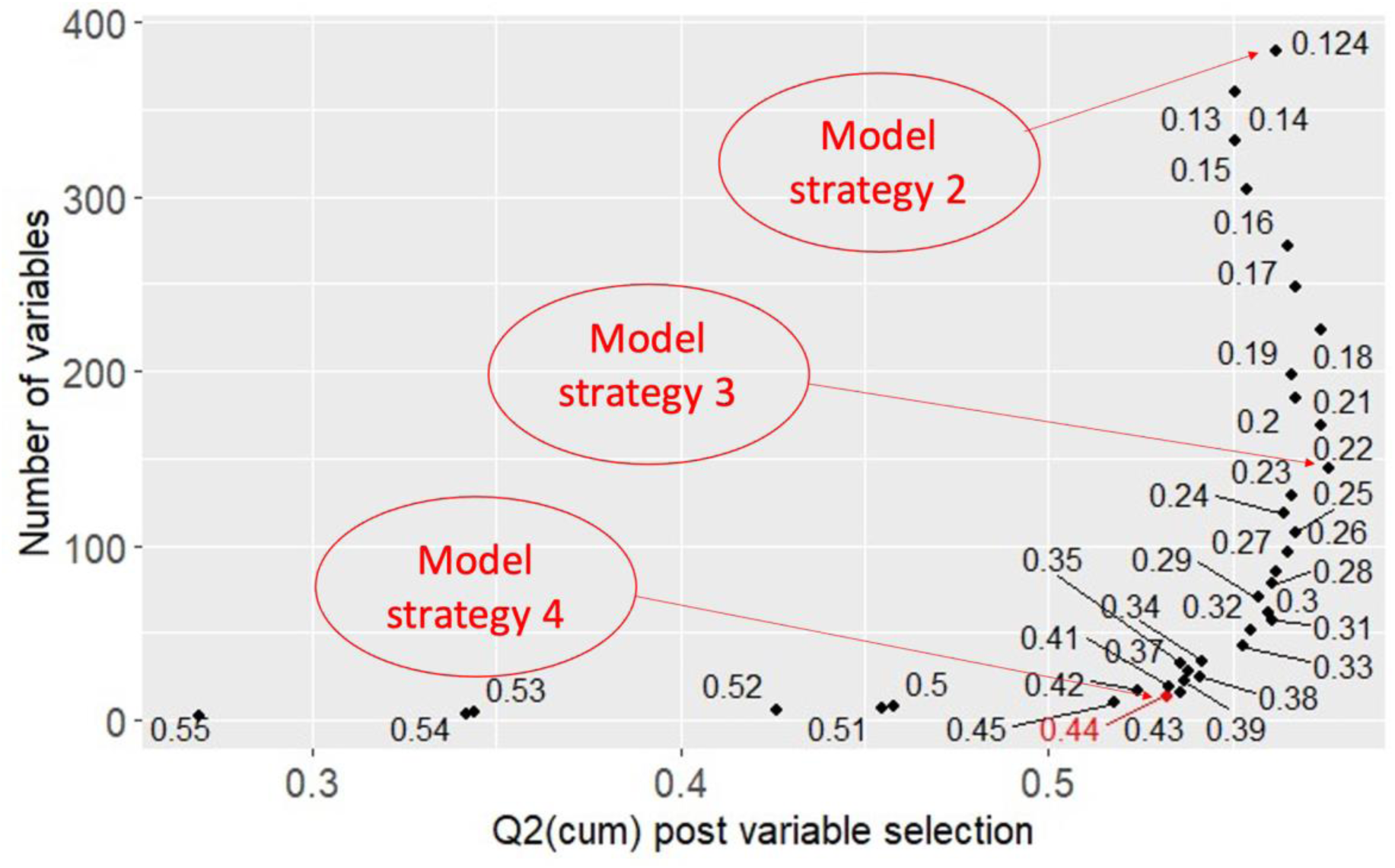
Visualization of the optimization of p(corr) cutoffs for different modeling strategies, for the comparison of E+P versus Nonuser, age 61-65 using Modeling strategy 2-4.

Figure 6 presents the optimized OPLS model for the comparison between E+P users and hormone non-users in the 61-65 age group, employing Modeling strategy 4. The separation between the two groups is shown by the score plot of the predictive component (Figure 6A). Additionally, the corresponding permutation over variable selection demonstrated high significance for both R^2^ and Q^2^, as indicated by p[R2_perm_over_vs], p[Q2_perm_over_vs], and the correlation coefficient between Y and permuted Y (Figure 6C-D).

**Figure 6.**
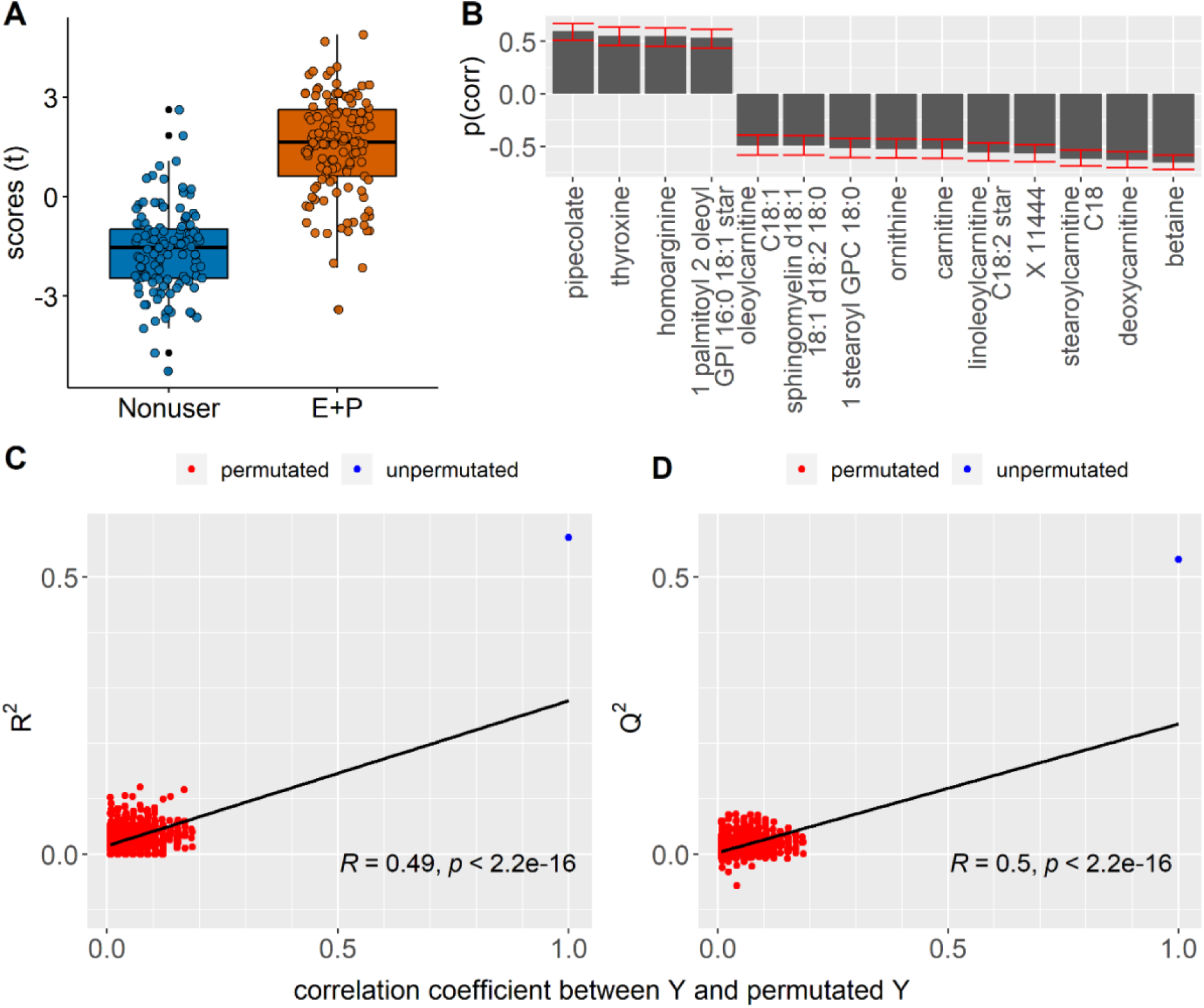
Scores plot (A) and the scaled loadings (B) of E+P users versus hormone Nonusers age group 61-65 years using Modelling strategy 4. Error bars in B indicate the 95% confidence interval of p(corr). Permutation over variable selection for the model is shown for R^2^ (C) and Q^2^ (D), with permuted models displayed in red and unpermuted models in blue. panels C-D: Displays R and p-value for Pearson’s correlation.

### Effect of Small Sample Sizes on R^2^ and Q^2^ and the Significance of Using Permutations

The study examined how decreasing sample size affects R^2^ and Q^2^, as well as its implications for significance using permutations pre-, post-, and over variable selection. Specifically, the robustness of a strong model comparing group E+P versus non-users was evaluated against a weak model of E-only versus non-users and a comparison expected to yield no significant models of non-users versus non-users, all within the 61-65 age group. Equal sample sizes were drawn from each group, ranging from n=4 to n=80, and constructed OPLS models using Modeling strategy 3.

With decreasing sample sizes, Q^2^ inflated, approaching 1.0 in models post-variable selection for both the strong and weak models, as well as when comparing non-users to non-users (Figure 8). The trend was similar for R^2^ (data not shown). In models pre-variable selection, decreasing sample size resulted in decreasing Q^2^ in the strong model (Figure S4-L) but was unaffected in the weak model (Figure S4-K) and increased when modeling non-users versus non-users (Figure S4-J).

When model significance was assessed using permutation post-variable selection, Q^2^ was significant for all models, including non-users versus non-users, regardless of sample size (Figure S4-D-F).

Examining the significance of Q^2^ using permutations over variable selection, the majority of the strong models remained significant with decreasing sample size (Figure S4-I), while half of the weak models were significant using Modeling strategy 3, with more significance at larger sample sizes (Figure S4-H). Thirteen percent of models comparing non-users to non-users were significant (Figure S4-G). When considering the regression coefficient (>0.1) and p-value for the regression coefficient over the permutations (<0.05), 6% of the models comparing non-users versus non-users were significant (Figure S4-J).

Assessing the significance of Q^2^ by permutating the models pre-variable selection, the strong models remained significant as long as more than 10 subjects were used in each group. As expected, almost all models comparing non-users versus non-users (i.e., no model) remained insignificant (Figure S4-J).

### Establishing a Statistically Sound Cutoff for Q^2^

To establish a significance level for Q^2^, the overinflation of the models was estimated by comparing the Q^2^ of models created from permuted data, including variable selection, to the Q^2^ of models without variable selection. Additionally, the difference between the Q^2^ of unpermuted models pre- and post-variable selection was estimated. These two estimates of overinflation overlapped each other at sample sizes below 20 in this dataset (Figure 7).

**Figure 7.**
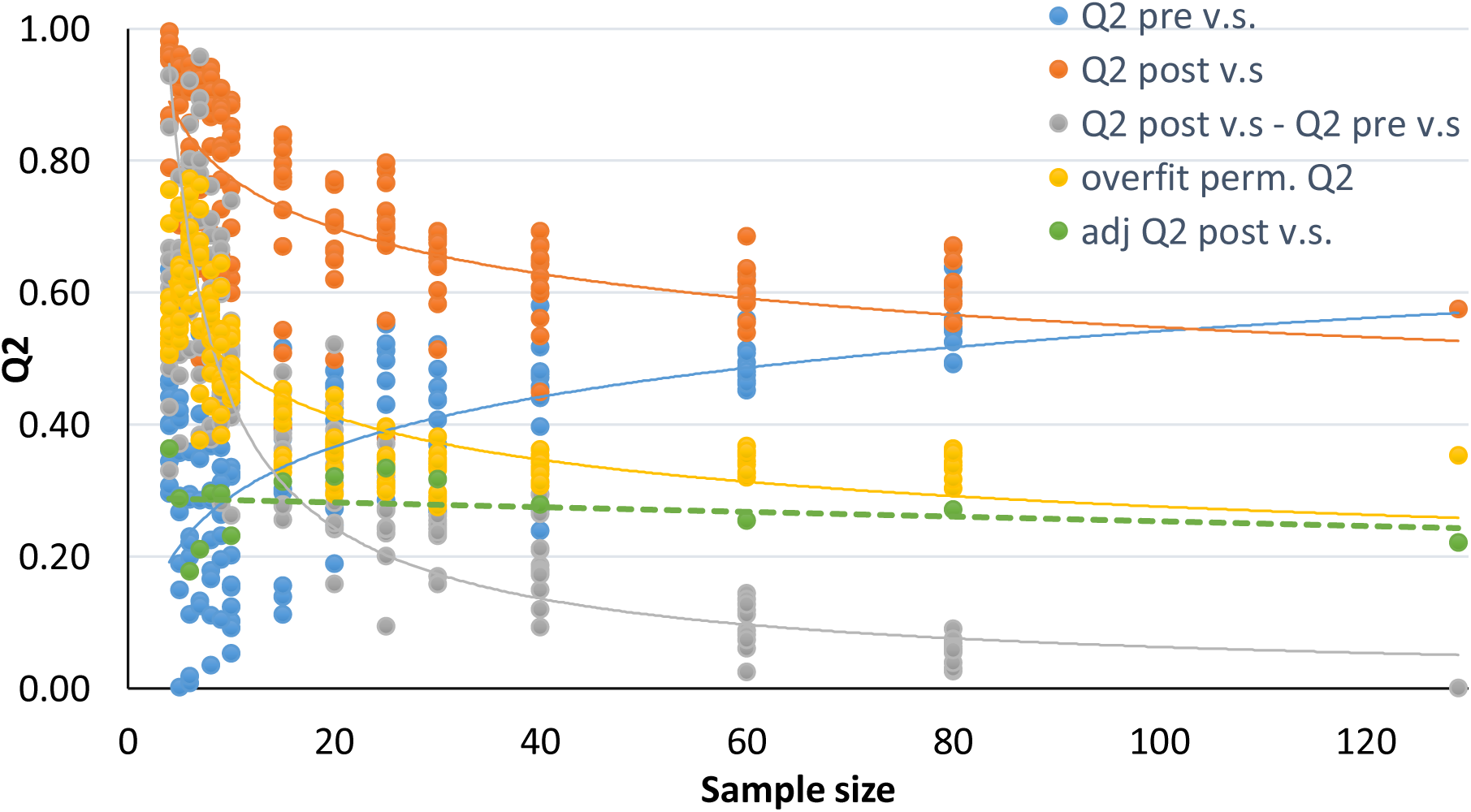
The figure displays Q^2^ versus group size of models comparing the groups of E+P versus Nonuser, age group 61-65. Q^2^ pre v.s: Q^2^ of models without prior variable selection; Q^2^ post v.s.: Q^2^ of models post variable selection; Q^2^ post v.s. – Q^2^ pre v.s.: The difference between Q^2^ pre and post variable selection; perm.overfit: Permuted overinflation is calculated by subtracting the average of the permuted models pre variable selection from the Q^2^ of the permuted models over variable selection; adj Q^2^ post v.s.: Adjusted Q^2^ post variable selection is obtained by subtracting the overinflation from the Q^2^ post variable selection.

An adjusted Q^2^ by subtracting the overinflation derived from permuted data from the Q^2^ of the models post-variable selection was calculated. This adjusted Q^2^ was independent of sample size and served as a more valid measure of how well a model performs than Q^2^ post-variable selection. A Q^2^ greater than zero indicates true variation larger than random explained by the model.

The number of orthogonal components was either decreased or not affected by decreasing sample sizes, indicating that the increase in R^2^ and Q^2^ was not due to the number of orthogonals.

## Discussion

Here, the widespread utilization of OPLS modeling in various omics fields for discerning group separation significance and identifying variables driving such separation is underscored. Despite its advantages over univariate statistics, OPLS modeling faces challenges, notably overfitting, particularly in small cohorts. The determination of significance thresholds for model statistics lacks clear guidelines, posing complexity and uncertainty, especially for the novice user. Cross-validation (CV), particularly Q^2^, serves as a gold standard for validation in small cohorts, but the applicable significance threshold remains elusive.

Lindgren et al. (1996) demonstrated the utility of permutations for OPLS models, showing that permutations over variable selection and pre-variable selection could distinguish a model from a randomly created one [11]. The authors also proposed estimating model overinflation by comparing Q^2^ of the model fitted on permuted data pre-variable selection to Q^2^ of the model fitted post-variable selection on the permuted data. In these studies, the effectiveness of permutation procedures with small sample sizes was tested, revealing that permutations pre- and over variable selection efficiently establish significance levels, even with smaller sample sizes. Specifically, OPLS modeling performance was examined on small sample sizes using a publicly available dataset and evaluated various permutation testing methods for assessing associated risks of overfitting or overinflation of model statistics. Our findings reveal a tendency of R^2^ and Q^2^ to increase with decreasing sample sizes. The results demonstrate that permutations pre- and over variable selection largely guard against the identification of random models as significant, while permutations post-variable selection exhibit limited efficacy, particularly with small samples. Additionally, the magnitude of Q^2^ overestimation resulting from variable selection was estimated, leading to an adjusted Q^2^ more resilient against group size differences, thus serving as a dataset-specific significance threshold.

Furthermore, ropls-ViPerSNet was introduced, an R workflow incorporating variable selection and permutations pre-, post-, and over variable selection, along with estimating overinflation and adjusted Q^2^. Its utility was demonstrated on the aforementioned publicly available metabolomic dataset, showcasing its automated comparison of pairwise cohort comparisons and stratified analyses. While the dataset’s lack of smoking status information limits its variance assessment, the script’s automation streamlines analysis, reduces subjectivity, and facilitates efficient reruns with new data or settings.

Another method suggested for testing overfitting or overinflation is the use of cross-validated score plots [36]. While it provides a more predictive picture of the score plot, it lacks a significance threshold. Future improvements to ropls-ViPerSNet could incorporate non-cross-validated score plots to address potential over-optimism.

Although ROC analysis is implemented in ropls-ViPerSNet using precrec and pROC packages, there’s potential for further enhancement by integrating AUROC into significance testing through permutations to establish a threshold for significance [37].

The ropls-ViPerSNet workflow is designed for ease in modeling numerous group comparisons and assessing significance using stringent permutation testing, particularly beneficial for screening many groups for differences. It offers flexibility and automation, especially suited for studying ropls model performance, particularly in scenarios with small sample sizes prone to overinflated model statistics. By rigorously testing for significance, the workflow contributes to reducing the high rate of false findings in scientific research, thereby enhancing the robustness of OPLS models.

Modeling strategies provided by the script offer versatility for various analysis goals, from pathway enrichment to biomarker identification. The flexibility and user-friendliness of ropls-ViPerSNet, alongside its potential for parallel computing, ensure applicability across omics platforms and clinical data analysis. Overall, this study highlights the importance of robust statistical approaches and automated workflows in enhancing the reliability and efficiency of OPLS modeling in metabolomics and related fields.

## Conclusion

The ropls-ViPerSNet workflow actively performs OPLS-DA modeling by semi-automatically optimizing models through variable selection. It also conducts permutation testing pre-, post-, and over variable selection to establish a significance level for the predictive model statistics Q^2^ of models post-variable selection.

Through extensive quality control, permutation testing is conducted pre-variable selection and over variable selection to mitigate the risk of false positives. Permutation pre-variable selection ensures significant differences between groups, while permutations over variable selection ensure that models maintain significance post-variable selection.

To demonstrate the utility of the workflow, it was applied to a publicly available metabolomic dataset. Our investigation into the effects of model on small group sizes reveals that R^2^ and Q^2^ increase with decreasing sample sizes. Both permutations pre-variable selection and permutations over variable selection are shown to significantly mitigate the identification of random models as significant, whereas permutations post-variable selection do not.

The proposed adjusted Q^2^, derived by subtracting the overinflation from the model post-variable selection, remains stable across sample sizes and may serve as a threshold for the significance of OPLS models. This is essential as traditional statistical evaluation alone may not adequately discern whether a model is significantly better than a random model post-variable selection.

---

**Supplementary Table 1.**
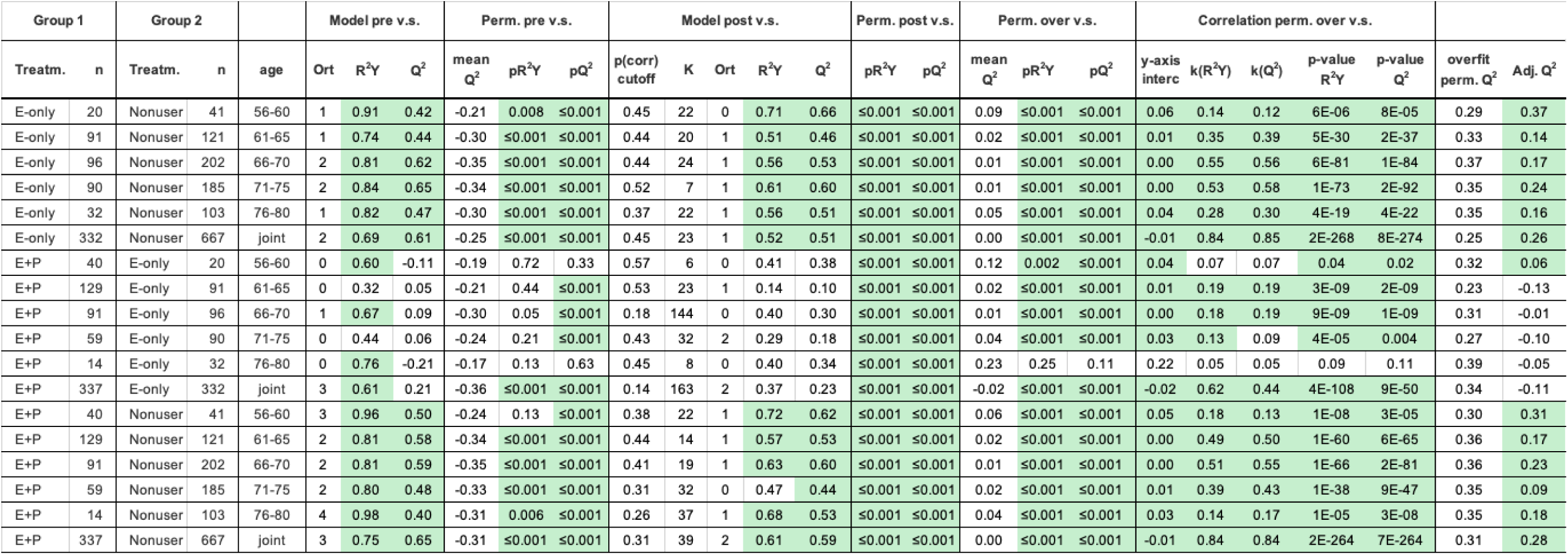
v.s.: variable selection; Perm: permutation analysis; Treatm: treatment group; n: number of subjects/group; Ort: number of orthongonal components in OPLS model; pR^2^Y: significance of nonparametric permutation analysis for R^2^Y; pQ^2^: significance of nonparametric permutation analysis for Q^2^; K: number of selected variables in optimized model; mean Q^2^: mean Q^2^ for 1000 permutations over v.s.; k(R^2^Y): Pearson correlation coefficient between R^2^Y and correlation between permuted and unpermuted group belonging; k(Q^2^): Pearson correlation coefficient between Q^2^ and correlation between permuted and unpermuted group belonging; p-value R^2^Y: significance of Pearson correlation coefficient between R^2^Y and correlation between permuted and unpermuted group belonging; p-value Q^2^: significance of Pearson correlation coefficient between Q^2^ and correlation between permuted and unpermuted group belonging; overfit perm Q^2^: mean (Q^2^ permuted models over v.s.)-mean (Q^2^ permuted models pre v.s.), represents an estimation of the overinflation occurring due to variable selection; Interc: intercept; Adj. Q^2^: Q^2^ post v.s. - overfit perm. Q^2^, represents estimation of Q^2^ corrected for overinflation by variable selection; E-only: group receiving estrogen hormone replacement therapy; Nonuser: group not receiving hormone replacement therapy; E+P: group receiving combined estrogen and progestin hormone replacement therapy.

**Supplementary Table 2.** Model statistics for OPLS models using Elastic Net variable selection.

| Dataset | R<sup>2</sup> | Q<sup>2</sup> | R<sup>2</sup>-Q<sup>2</sup> | # Pred | #ortho | pR<sup>2</sup>Y | pQ<sup>2</sup> |
| --- | --- | --- | --- | --- | --- | --- | --- |
| E_none_56_60_ortho_0 | 0.78 | 0.75 | 0.03 | 1 | 0 | <0.01 | <0.01 |
| E_none_56_60_ortho_1 | 0.84 | 0.81 | 0.03 | 1 | 1 | <0.01 | <0.01 |
| E_none_56_60_ortho_2 | 0.86 | 0.82 | 0.04 | 1 | 2 | <0.01 | <0.01 |
| E_none_61_65_ortho_0 | 0.46 | 0.45 | 0.01 | 1 | 0 | <0.01 | <0.01 |
| E_none_61_65_ortho_1 | 0.67 | 0.64 | 0.03 | 1 | 1 | <0.01 | <0.01 |
| E_none_61_65_ortho_2 | 0.69 | 0.64 | 0.05 | 1 | 2 | <0.01 | <0.01 |
| E_none_66_70_ortho_0 | 0.52 | 0.48 | 0.04 | 1 | 0 | <0.01 | <0.01 |
| E_none_66_70_ortho_1 | 0.77 | 0.72 | 0.05 | 1 | 1 | <0.01 | <0.01 |
| E_none_66_70_ortho_2 | 0.84 | 0.76 | 0.08 | 1 | 2 | <0.01 | <0.01 |
| E_none_71_75_ortho_0 | 0.72 | 0.67 | 0.05 | 1 | 0 | <0.01 | <0.01 |
| E_none_71_75_ortho_1 | 0.83 | 0.77 | 0.06 | 1 | 1 | <0.01 | <0.01 |
| E_none_71_75_ortho_2 | 0.86 | 0.80 | 0.06 | 1 | 2 | <0.01 | <0.01 |
| E_none_76_80_ortho_0 | 0.76 | 0.60 | 0.15 | 1 | 0 | <0.01 | <0.01 |
| E_none_76_80_ortho_1 | 0.89 | 0.78 | 0.11 | 1 | 1 | <0.01 | <0.01 |
| E_none_76_80_ortho_2 | 0.94 | 0.85 | 0.09 | 1 | 2 | <0.01 | <0.01 |
| E_none_all_ortho_0 | 0.57 | 0.54 | 0.03 | 1 | 0 | <0.01 | <0.01 |
| E_none_all_ortho_1 | 0.68 | 0.65 | 0.03 | 1 | 1 | <0.01 | <0.01 |
| E_none_all_ortho_2 | 0.77 | 0.71 | 0.06 | 1 | 2 | <0.01 | <0.01 |
| E_P_E_61_65_ortho_0 | 0.72 | 0.42 | 0.29 | 1 | 0 | <0.01 | <0.01 |
| E_P_E_61_65_ortho_1 | 0.82 | 0.59 | 0.23 | 1 | 1 | <0.01 | <0.01 |
| E_P_E_61_65_ortho_2 | 0.90 | 0.63 | 0.27 | 1 | 2 | <0.01 | <0.01 |
| E_P_E_66_70_ortho_0 | 0.66 | 0.57 | 0.09 | 1 | 0 | <0.01 | <0.01 |
| E_P_E_66_70_ortho_1 | 0.73 | 0.61 | 0.13 | 1 | 1 | <0.01 | <0.01 |
| E_P_E_66_70_ortho_2 | 0.76 | 0.59 | 0.17 | 1 | 2 | <0.01 | <0.01 |
| E_P_E_71_75_ortho_0 | 0.80 | 0.68 | 0.12 | 1 | 0 | <0.01 | <0.01 |
| E_P_E_71_75_ortho_1 | 0.87 | 0.72 | 0.16 | 1 | 1 | <0.01 | <0.01 |
| E_P_E_71_75_ortho_2 | 0.90 | 0.71 | 0.19 | 1 | 2 | <0.01 | <0.01 |
| E_P_E_76_80_ortho_0 | 0.87 | 0.74 | 0.13 | 1 | 0 | <0.01 | <0.01 |
| E_P_E_76_80_ortho_1 | 0.93 | 0.80 | 0.12 | 1 | 1 | <0.01 | <0.01 |
| E_P_E_76_80_ortho_2 | 0.95 | 0.80 | 0.15 | 1 | 2 | <0.01 | <0.01 |
| E_P_E_all_ortho_0 | 0.42 | 0.26 | 0.17 | 1 | 0 | <0.01 | <0.01 |
| E_P_E_all_ortho_1 | 0.56 | 0.32 | 0.23 | 1 | 1 | <0.01 | <0.01 |
| E_P_E_all_ortho_2 | 0.66 | 0.40 | 0.25 | 1 | 2 | <0.01 | <0.01 |
| E_P_none_56_60_ortho_0 | 0.85 | 0.81 | 0.05 | 1 | 0 | <0.01 | <0.01 |
| E_P_none_56_60_ortho_1 | 0.95 | 0.89 | 0.06 | 1 | 1 | <0.01 | <0.01 |
| E_P_none_56_60_ortho_2 | 0.97 | 0.90 | 0.07 | 1 | 2 | <0.01 | <0.01 |
| E_P_none_61_65_ortho_0 | 0.52 | 0.51 | 0.01 | 1 | 0 | <0.01 | <0.01 |
| E_P_none_61_65_ortho_1 | 0.70 | 0.69 | 0.01 | 1 | 1 | <0.01 | <0.01 |
| E_P_none_61_65_ortho_2 | 0.70 | 0.69 | 0.01 | 1 | 2 | <0.01 | <0.01 |
| E_P_none_66_70_ortho_0 | 0.65 | 0.56 | 0.08 | 1 | 0 | <0.01 | <0.01 |
| E_P_none_66_70_ortho_1 | 0.84 | 0.77 | 0.07 | 1 | 1 | <0.01 | <0.01 |
| E_P_none_66_70_ortho_2 | 0.89 | 0.81 | 0.09 | 1 | 2 | <0.01 | <0.01 |
| E_P_none_71_75_ortho_0 | 0.76 | 0.69 | 0.07 | 1 | 0 | <0.01 | <0.01 |
| E_P_none_71_75_ortho_1 | 0.83 | 0.74 | 0.09 | 1 | 1 | <0.01 | <0.01 |
| E_P_none_71_75_ortho_2 | 0.84 | 0.75 | 0.09 | 1 | 2 | <0.01 | <0.01 |
| E_P_none_76_80_ortho_0 | 0.76 | 0.65 | 0.11 | 1 | 0 | <0.01 | <0.01 |
| E_P_none_76_80_ortho_1 | 0.81 | 0.72 | 0.09 | 1 | 1 | <0.01 | <0.01 |
| E_P_none_76_80_ortho_2 | 0.82 | 0.73 | 0.09 | 1 | 2 | <0.01 | <0.01 |
| E_P_none_all_ortho_0 | 0.56 | 0.52 | 0.03 | 1 | 0 | <0.01 | <0.01 |
| E_P_none_all_ortho_1 | 0.74 | 0.70 | 0.04 | 1 | 1 | <0.01 | <0.01 |
| E_P_none_all_ortho_2 | 0.81 | 0.75 | 0.06 | 1 | 2 | <0.01 | <0.01 |
| E_P_none_61_65_ortho_0_125 | 0.66 | 0.63 | 0.04 | 1 | 0 | <0.01 | <0.01 |
| E_P_none_61_65_ortho_0_250 | 0.52 | 0.51 | 0.01 | 1 | 0 | <0.01 | <0.01 |
| E_P_none_61_65_ortho_0_40 | 0.77 | 0.75 | 0.02 | 1 | 0 | <0.01 | <0.01 |
| E_P_none_61_65_ortho_0_50 | 0.76 | 0.76 | 0.01 | 1 | 0 | <0.01 | <0.01 |
| E_P_none_61_65_ortho_0_65 | 0.86 | 0.79 | 0.07 | 1 | 0 | <0.01 | <0.01 |
| E_P_none_61_65_ortho_1_125 | 0.70 | 0.67 | 0.03 | 1 | 1 | <0.01 | <0.01 |
| E_P_none_61_65_ortho_1_250 | 0.70 | 0.68 | 0.02 | 1 | 1 | <0.01 | <0.01 |
| E_P_none_61_65_ortho_1_40 | 0.79 | 0.74 | 0.05 | 1 | 1 | <0.01 | <0.01 |
| E_P_none_61_65_ortho_1_50 | 0.87 | 0.83 | 0.05 | 1 | 1 | <0.01 | <0.01 |
| E_P_none_61_65_ortho_1_65 | 0.96 | 0.88 | 0.08 | 1 | 1 | <0.01 | <0.01 |
| E_P_none_61_65_ortho_2_125 | 0.70 | 0.68 | 0.03 | 1 | 2 | <0.01 | <0.01 |
| E_P_none_61_65_ortho_2_250 | 0.70 | 0.67 | 0.02 | 1 | 2 | <0.01 | <0.01 |
| E_P_none_61_65_ortho_2_50 | 0.87 | 0.83 | 0.05 | 1 | 2 | <0.01 | <0.01 |
| E_P_none_61_65_ortho_2_65 | 0.98 | 0.90 | 0.08 | 1 | 2 | <0.01 | <0.01 |

**Figure S1.**
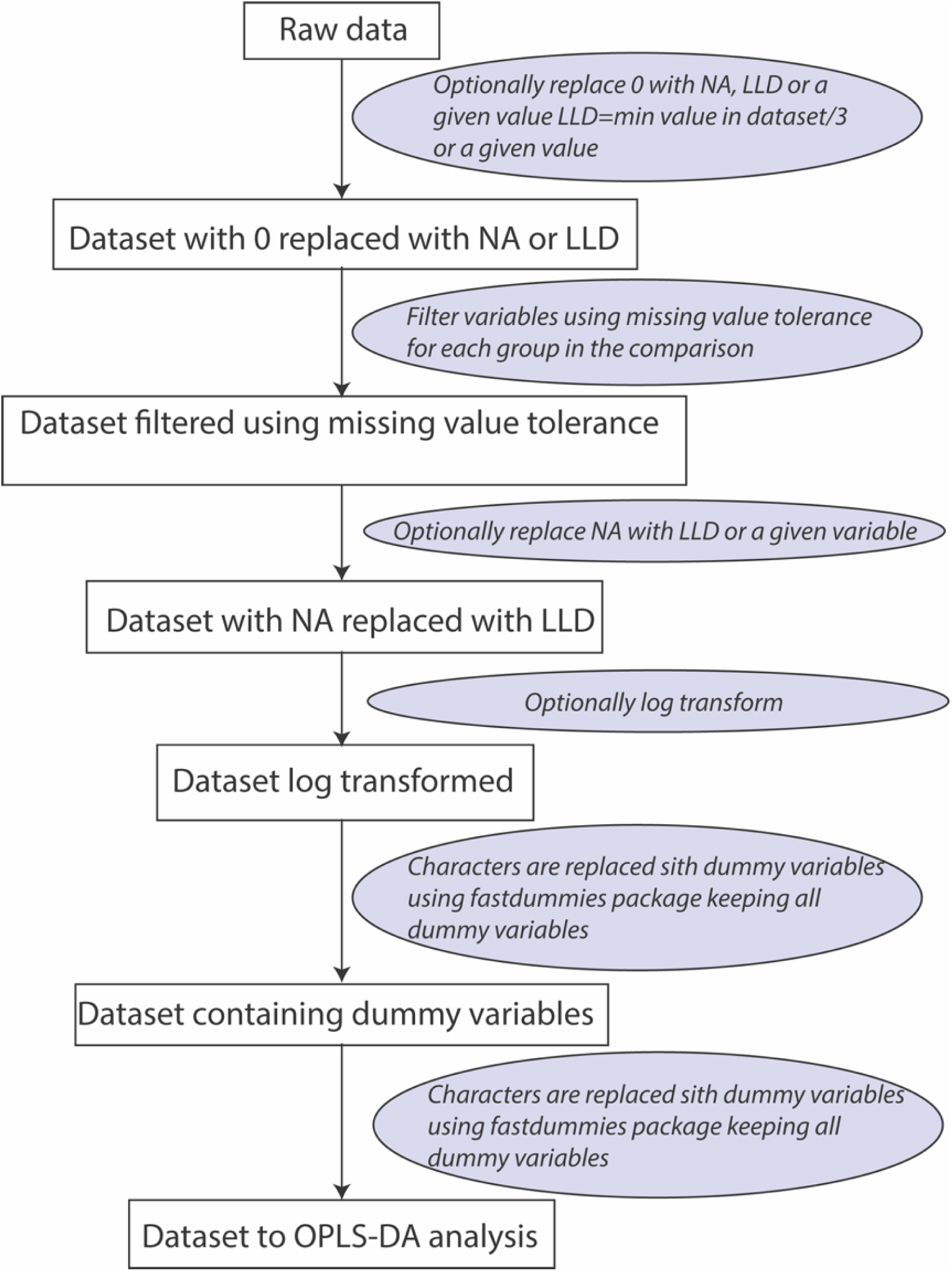
Schematic of data pre-processing options in the ropls-ViPerSNet workflow.

**Figure S2.**
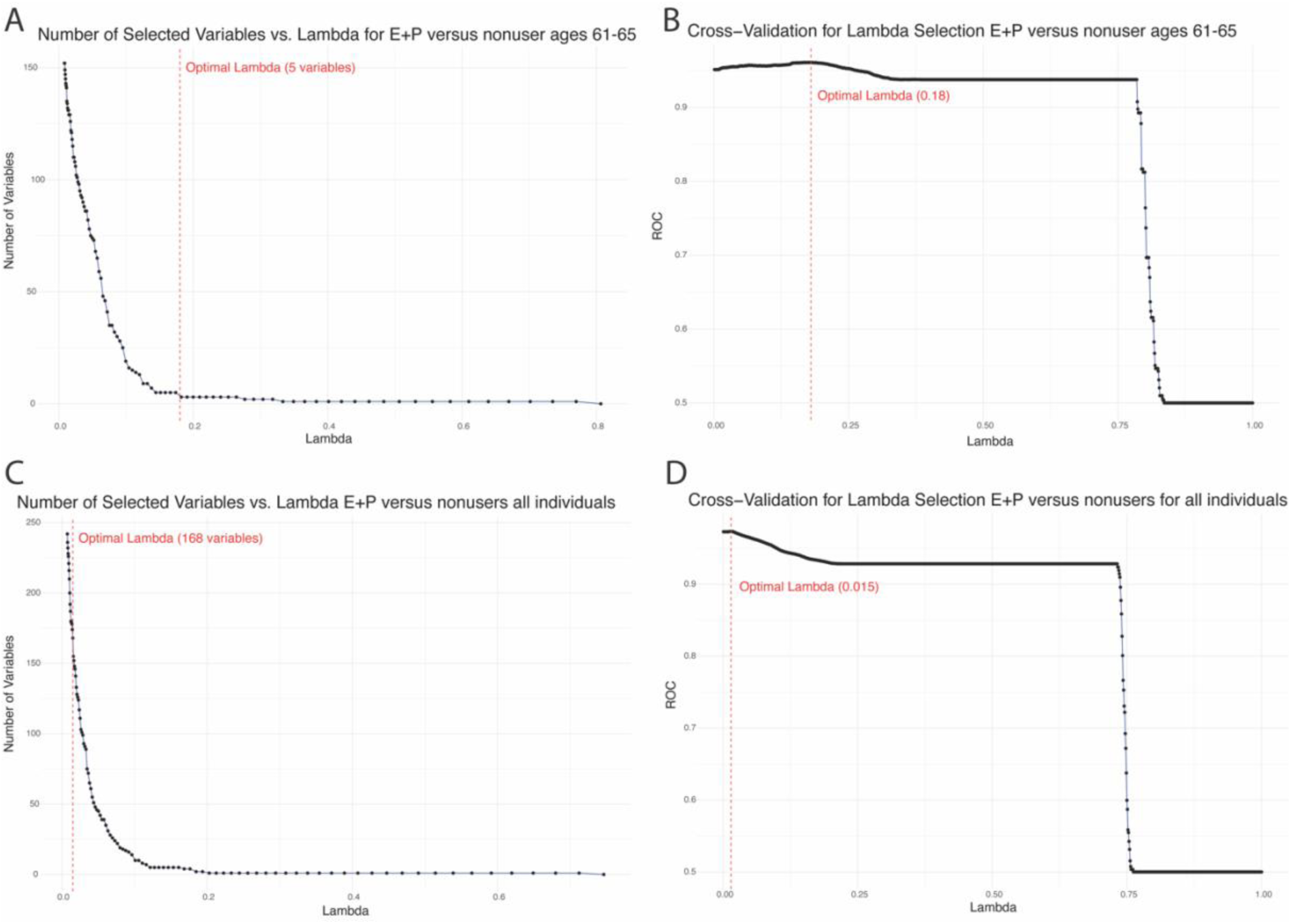
Optimization of Lambda for the model and variables. These figures are based on E+P versus nonusers for all individuals (C and D) and for age stratified group 61-65 years (A and B).

**Figure S3.**
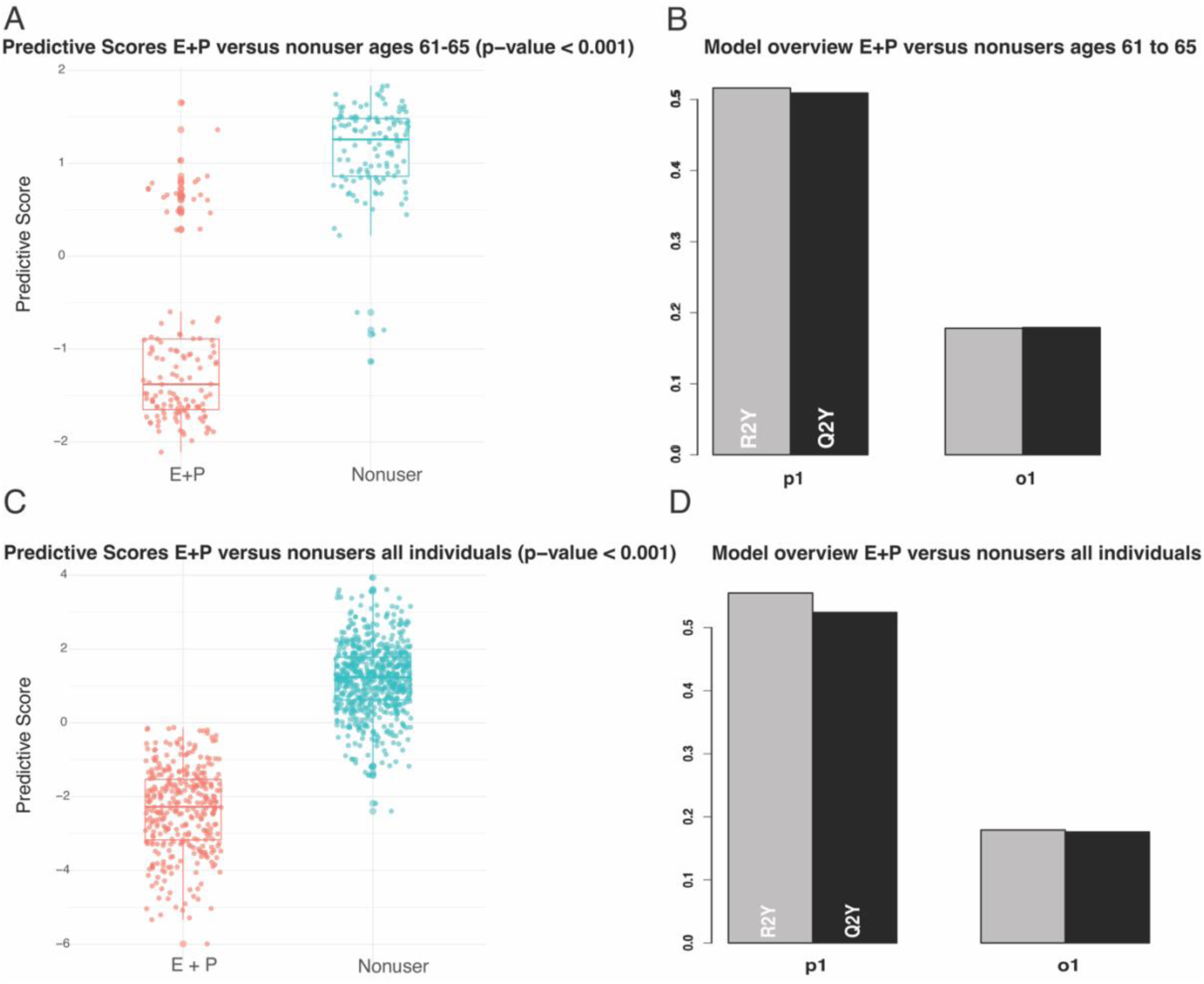
Results from the Eastic Net narrowed OPLS model for E+P versus nonusers based on one predictive and one orthogonal component. Both models were significant. Q^2^ slightly improved when the analysis was performed on age stratified group 61-65.

**Figure S4:**
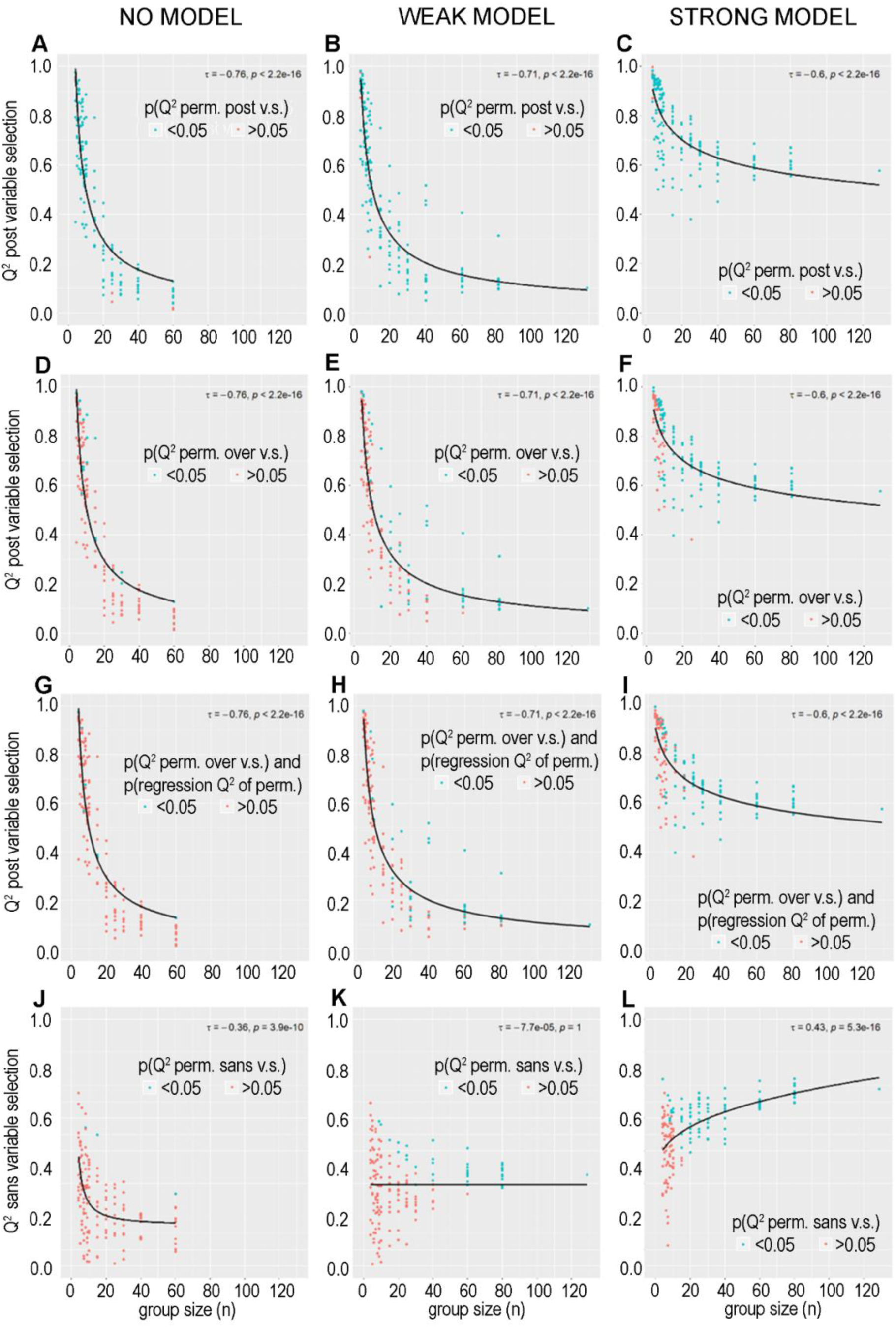
Q^2^ of models post variable selection using Modelling strategy 3 was significant across all samples sizes and models (A-C, colored by p[Q^2^ permuted post variable selection]<0.05). However, the majority of the models in the weak- and non-models did not pass permutation over variable selection (D-E), whereas the majority of the strong models passed permutation over variable selection at a cutoff of n=8 (panel F). The same trend held true when also including the significance of the correlation of the permutation, but reduced the false discovery rate to 6%(panels G-I). Permutation sans variable selection displayed the opposite trend, with a decreasing Q^2^ with decreasing group size in the strong model (panel L). This trend may either imply that this approach is too stringent, particularly at group sizes below n=10, or simply reflect the decreasing statistical power resulting from the smaller n. As expected, the non-models did not pass the sans v.s. permutations tests (J)-The weak model required a sample size of n=40-60 to identify the models as significant using permutation test (K).

